# Human medial pulvinar is involved in face and tactile processing and dynamically adjusts its cortical connectivity based on ongoing stimulation

**DOI:** 10.1101/2024.09.20.614101

**Authors:** M. Gaudet-Trafit, M. Froesel, S. Dali, F. Lamberton, S. Ben Hamed

## Abstract

Optimal processing of our surrounding environment relies on our ability to detect and integrate external information through multiple sensory modalities to build a coherent representation of the world. Interactions between cortical and subcortical structures contribute to this process. In this context, there is growing evidence that the pulvinar plays a crucial role in visual and face processing. In contrast, its role in auditory, voice and tactile processing has been poorly investigated. Here, we use fMRI localizer tasks to describe pulvinar functional organization and functional connectivity with the brain during face, voice and tactile processing. We reproduce the activation of the ventral part of the medial pulvinar in face perception and we describe an increased pulvinar connectivity with face processing areas and a decreased connectivity with low level visual areas during face processing. In addition, we describe activations of the medial pulvinar during air-puff face, hands and feet tactile stimulations and changes in pulvino-cortical connectivity as a function of which body part is being stimulated. No activations are observed during either voice or non-voice stimulations. Overall, this supports a role of medial pulvinar in multisensory processing and the modulation of cortical areas as a function of the sensory context.

## Introduction

Optimal processing of the surrounding environment and the identification of the most relevant incoming information is crucial for navigating and adapting in a complex dynamic environment. Sensory processing is carried out by both cortical and subcortical areas of the central nervous system (Mesulam, 1998; Murray and Wallace, 2012). The processing of social sensory signals involved in primate communication is mainly carried out by visual and auditory brain systems. In the visual system, the processing of faces is crucial to social cognition. A large network of regions spanning from the occipital to frontal cortices including the fusiform (FFA), occipital face area (OFA), posterior superior temporal lobe (pSTS, also called face-responsive STS, fSTS), anterior temporal lobe (ATL) and orbito-frontal cortex (OFC) has been associated with visual face processing (Gratton et al., 2013; Haxby et al., 2000; Schobert et al., 2018; Troiani et al., 2016; Zhang et al., 2009). In the auditory system, in both human and non-human primates, a voice patch system similar to the face patch system was recently described (Bodin and Belin, 2020; Pernet et al., 2015). Three main voice patches have been identified along the superior temporal gyrus and sulcus, a posterior voice patch, a median voice patch and an anterior voice patch with a large interindividual variability (Bodin and Belin, 2020).

In this context, it was initially proposed that subcortical regions such as the thalamus are mere sensory relays, while cortical areas are responsible for the implementation of higher order cognitive functions such as social cognitive functions. This view has been challenged over the last years. Notably, there is growing evidence that the pulvinar, the largest thalamic nucleus, plays a crucial role in higher cognitive functions and prioritizes incoming sensory information including social-related sensory information (Arcaro et al., 2018; Froesel et al., 2021; Wen et al., 2023). In addition to visual and audio-visual processing (Froesel et al., 2024a; Vittek et al., 2023) the pulvinar has been shown to be involved in sensorimotor processing (Charyasz et al., 2023). While somatotopic information from periphery to cortex, has been reliably described to be relayed by the ventro-posterior complex of the thalamus (Delhaye et al., 2018), in non-human primates, the anterior subdivision of the pulvinar has anatomic connections to parietal areas 1, 2 and 7b, lateral parietal cortex (Delhaye et al., 2018) and has been shown to be responsive to puff stimulations on the face as well as to visuo-tactile stimulation ( Froesel et al., 2021).

The pulvinar is a posterior thalamic nucleus covering 30 % of the thalamus. It exhibits strong interconnexions with the primary and associative cortices (Froesel et al., 2021; Shipp, 2003). It has a complex and rich structural and functional connectivity organization with the cortex (Asanuma et al., 1985; Froesel et al., 2021; Froesel et al., 2024b; Leh et al., 2008; Shipp, 2003; Vittek et al., 2023). This extensive connectivity has led this nucleus to be considered as a cortical hub for various cognitive processes such as attention, salience or social cognition (Benarroch, 2015; Saalmann and Kastner, 2011; Froesel et al 2021). These studies highlight the role of pulvinar as a coordinator and modulator of perception primarily focusing on vision (Benarroch, 2015; Fiebelkorn et al., 2019; Saalmann and Kastner, 2011). In particular, dorsal pulvinar has been associated with attention while ventral pulvinar has been associated visual temporal structure processing and spatial position and categorization in the context of visual face and scene processing. Accordingly, these subdivisions are functionally connected with specific cortical areas (Arcaro et al., 2018). In primates, the pulvinar is classically divided in several sub-regions based on their specific cytoarchitectonic and chemoarchitectonic properties, namely the anterior, medial, lateral and inferior subdivisions, each containing further subdivisions (Gutierrez et al., 2000, 1995; Morel et al., 1997; Olszewski, 1952; Stepniewska et al., 1999; Walker, 1938). Aside from this anatomical description of the pulvinar, a meta-analysis of task-related neuroimaging studies identified five pulvinar clusters: inferior, lateral, medial, anterior and superior pulvinar cluster (Barron et al., 2015). In contrast, pulvinar clustering based on functional connectivity studies splits the pulvinar into a dorso-medial, a ventro-medial, a lateral, an anterior and an inferior cluster (Guedj and Vuilleumier, 2020). Thus, the parcellation of this nucleus is challenging, three different methods resulting in three different parcellations.

Although the role of the pulvinar in vision processing has been well studied, its involvement in auditory and tactile processing remains less understood. The exploration of these sensory modalities is essential to comprehensively elucidate the pulvinar’s multifaceted functions. Additionally, investigating the functional connectivity patterns of the pulvinar across different sensory modalities and sensory contexts can provide deeper insights into how this thalamic structure integrates and coordinates the processing of sensory. In this study, we aim to investigate the functional role of pulvinar in face, voice and tactile processing and elucidate the functional connectivity of the responsive pulvinar regions of interest (seeds) with the rest of the brain. Participants performed three fMRI localizer tasks. First, a face localizer task in which participants were exposed to human faces, objects and scenes. Second, a voice localizer task in which participants were exposed to human voices and different non-voice sounds. Third, a tactile localizer task in which the face (cheeks), the hands and the feet of participants were stimulated unilaterally by air puffs. Our results revealed a localized activation of medial pulvinar in both face and tactile tasks but no response during the auditory task. Specifically, the central part of medial pulvinar was involved in face processing and medial regions were involved in tactile processing. We further characterize the dynamic connectivity patterns of pulvinar with the cortical domain as a function of the ongoing sensory stimulation. In particular, we identify specific pulvino-cortical connectivity patterns for visual face processing as well as for tactile processing.

## Results

The main purpose of this study was to explore the role of pulvinar in face, voice and tactile processing. In a first step, we validated our localizer tasks (figure 1) by describing the cortical and subcortical network that they activated. In a second step, we investigated pulvinar activations during the face, the voice and the tactile localizer tasks performing group-level statistical analyses on whole brain BOLD activity. In a last step, we used the pulvinar activations as seeds for a seed-based analysis, in order to characterize their functional connectivity patterns with the rest of the brain both during resting state and during the different localizer tasks.

**Figure 1:**
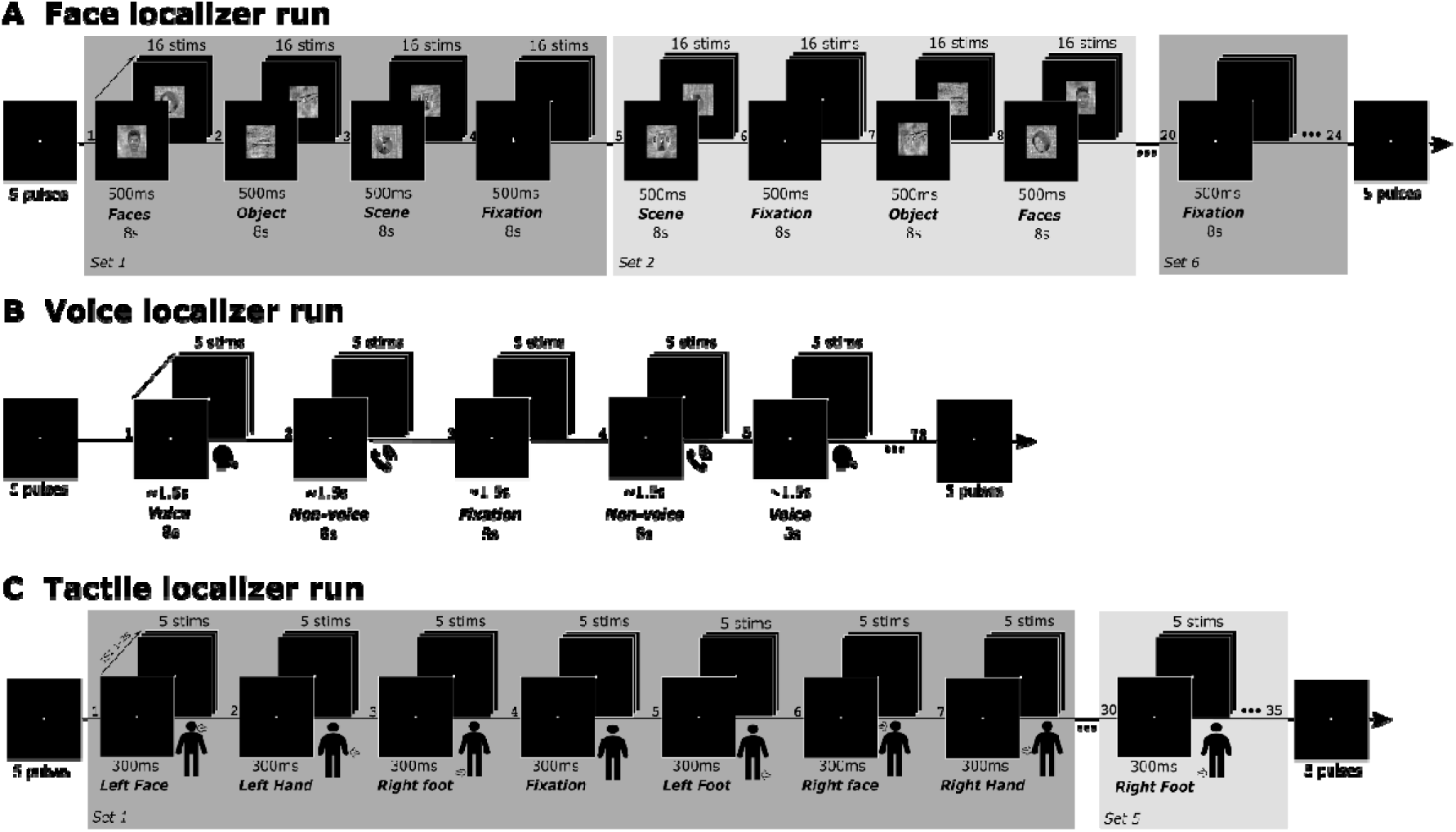
Face, tactile and voice localizer tasks. A) Face localizer. Each run was composed of 6 sets of 4 blocks (total of 24 blocks). Each set was composed of one face block, one object block, one scene block, and one fixation block presented in a pseudo-randomized order. Each block lasted 8s and involved the presentation of 16 randomized stimuli of 500ms. B) Voice Localizer. Each run was composed by 72 randomized blocks of voice stimuli, non-voice stimuli and fixation. Each block involved the repetition of 5 auditory stimuli of ∼1.5s separated by 15ms for a total duration of 8s. C) Tactile localizer. Each run was composed of 5 sets of 7 blocks each (total of 35 blocks). Each set was composed of a pseudorandomized presentation of left and right face, hand and feet blocks as well as one fixation blocks. Each block lasted 1.5s and involved the presentation of 5 repetitions of a 300ms stimulation.

### Face localizer cortical and subcortical activations

Two main contrasts were defined to analyze the activations during the Face localizer task: 1) a contrast comparing the face, object and scene conditions to fixation (face+object+scene>fixation a.k.a all>fixation); 2) a specific contrast comparing the face condition to the non-face conditions (face>object+scene). For the all>fixation contrast, cortical activations were located within a large cortical network encompassing bilateral occipital cortex, fusiform cortex and parietal cortex (figure 2, top panel and table S1.1). Subcortical activations were observed in the thalamus, superior colliculus, lateral and medial geniculate nuclei (figures 2 and 3B, and table S1.1) and hippocampus (figure 2 and table S1.1), bilaterally. For the face>object+scene contrast, we identified a large cortical activation cluster encompassing bilateral occipital cortex and fusiform cortex in addition to the ventral part of the right medial pulvinar and bilateral hippocampus. All activations for both contrasts were reported with FWE correction for multiple comparisons at voxel level p<0.05. Thus, the Face localizer robustly activated a cortical-subcortical network involved in visual processing. During face processing, a more restricted network is identified, coinciding with the classically reported face areas (Gratton et al., 2013; Haxby et al., 2000; Schwiedrzik et al., 2015; Tsao et al., 2008; Zhang et al., 2009).

**Figure 2:**
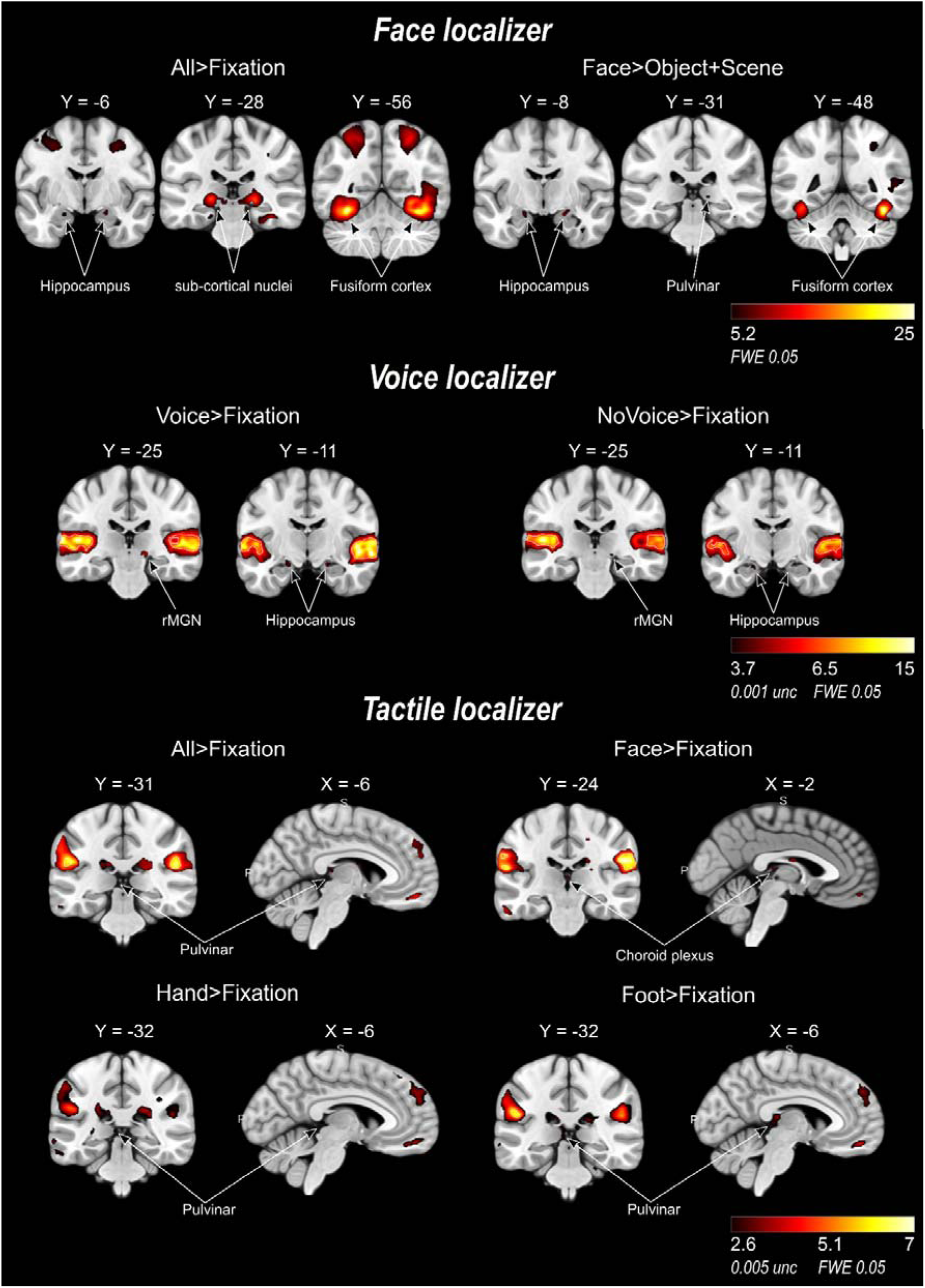
Whole brain contrasts activation for face, vocal and tactile localizer tasks. **(Top)** Face localizer. Coronal view sections of group level activation maps thresholded at p<0.05 FWE at voxel level and for all>fixation and face>object+scene contrasts. (Middle) Voice localizer. Coronal view sections of group level activation maps thresholded at p<0.05 FWE at voxel level (white contour on activation maps) and at p>0.001 uncorrected for voice>fixation and non-voice>fixation contrasts. (Bottom) Tactile localizer. Coronal view sections of group level activation maps thresholded at p<0.05 FWE at voxel level (white contour on activation maps) and at p>0.001 uncorrected for all>fixation, face>fixation, hand>fixation and foot>fixation contrasts. MNI coordinates are displayed above each section.

**Figure 3:**
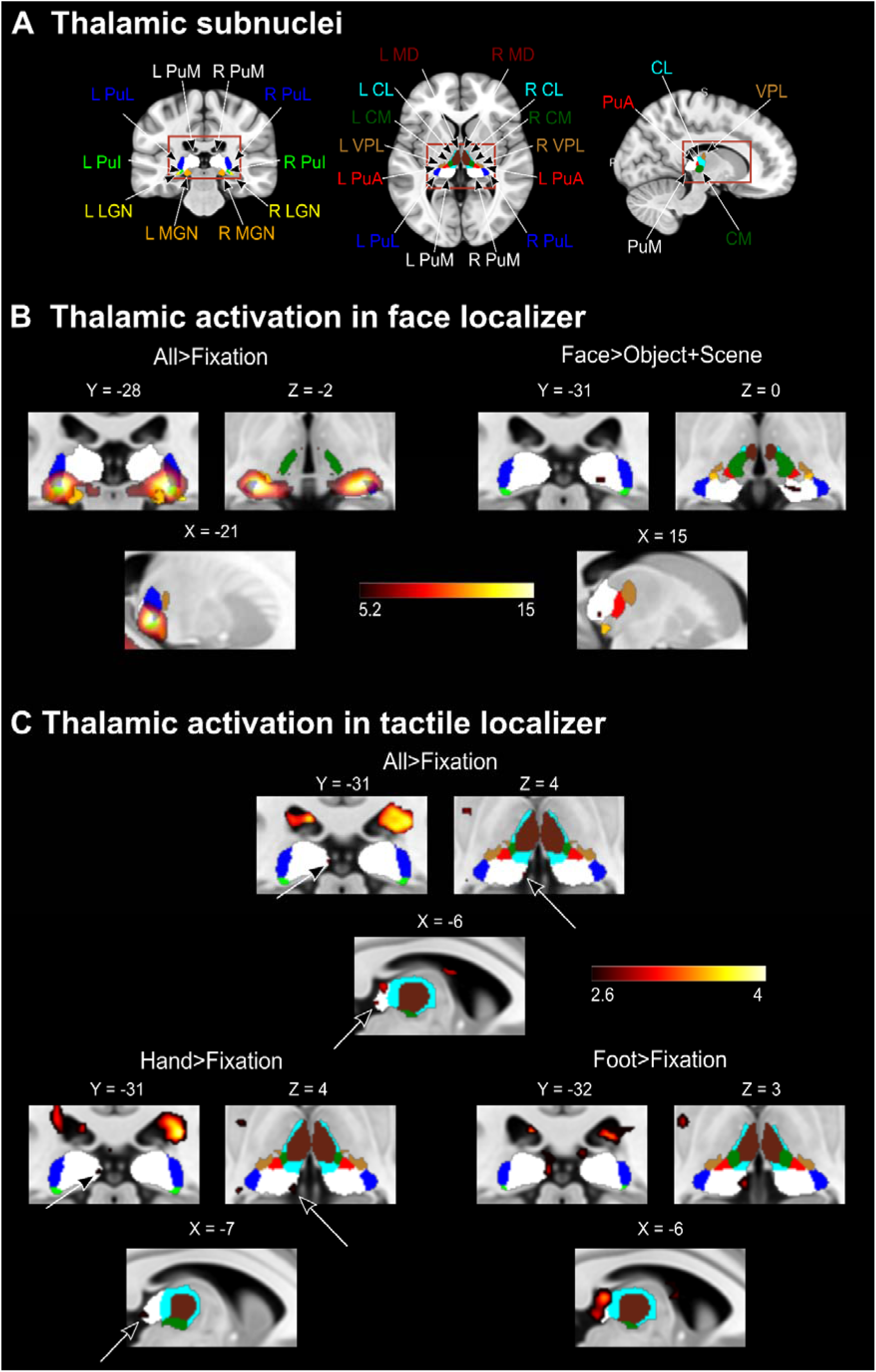
Thalamic activation in face and tactile localizer tasks. A) Thalamic subnuclei location based on the Morel human thalamic atlas (Morel et al., 1997). B) Face localizer. Coronal, axial and sagittal sections of group level activation thresholded maps (p<0.05 FWE at voxel level), for all>fixation and face>object+scene contrasts. C) Tactile localizer. Coronal, axial and sagittal sections of group level activation thresholded maps (p<0.005 uncorrected at voxel level), for all>fixation, hand>fixation and foot>fixation contrasts. Thalamic nuclei and subcortical structures and are shown in different colors: PuI: inferior pulvinar (green), PuA: anterior pulvinar (red), PuL: lateral pulvinar (blue), PuM: medial pulvinar (white), MD: mediodorsal nucleus (brown), CM: centre median nucleus (dark green), CL: central lateral nucleus (cyan), VPL: ventral posterior lateral nucleus (gold), MGN: medial geniculate nucleus (orange), LGN: lateral geniculate nucleus (yellow). MNI x, y, z coordinates are displayed above each section.

### Voice localizer cortical and subcortical activations

Three main contrasts were defined to analyze the activations during the Voice localizer task: 1) a voice>non-voice contrast; 2) a voice>fixation contrast; 3) a non-voice>fixation contrast. Fixation blocks were silent blocks in which no sound was displayed. In the voice>non-voice contrast, significant activations were observed within three cortical clusters in the left temporal mid gyrus, the right superior temporal gyrus and the right temporal mid gyrus (p<0.05, FWE correction for multiple comparison at voxel level, figure 2, Table S1.2). In the voice>fixation contrast, activations were observed mainly in the left and right temporal cortex (p<0.05, FWE correction for multiple comparison at voxel level, figure 2, Table S1.2). At uncorrected p<0.001 threshold at voxel level, we additionally identified activations in the right medial geniculate nucleus (MGN) and bilateral hippocampus (figure 2, Table S1.2). In the non-voice>fixation, two large clusters in the left and right temporal lobe were significantly activated at p<0.05, FWE correction for multiple comparison at voxel level (figures 2, Table S1.2). At uncorrected threshold p<0.001 at voxel level, activations were observed in the medial geniculate nucleus (MGN) and bilateral hippocampus (figures 2, Table S1.2). In summary, voice and no voice stimuli activated cortical temporal areas. Voices specifically activates the voice temporal patches. Right MGN and bilateral hippocampus were activated at lower threshold with no specificity for voice processing. No activations within the pulvinar nucleus were observed at these statistical thresholds. Thus, the Voice localizer robustly activated a cortical-subcortical network involved in auditory processing. During voice processing, a more restricted network is identified, coinciding with the classically reported voice areas (Belin et al., 2000; Bodin and Belin, 2020; Moerel et al., 2014; Pernet et al., 2015; Zhang et al., 2021).

### Tactile localizer cortical and subcortical activations

Four main contrasts were defined to analyze the activations during the Tactile localizer task: 1) a face>fixation contrast; 2) a hand>fixation contrast; 3) a foot>fixation contrast; 4) all stimulation conditions vs contrast, a.k.a. all>fixation contrast. For the all>fixation contrast, at a threshold of p<0.05 FWE corrected, we observed activations within the superior temporal gyrus, the supramarginal area and the Rolandic operculum bilaterally (figure 2, Table S1.3). We also observed activations in the left angular gyrus and in the ventral part of the left postcentral gyrus. At an uncorrected threshold of p<0.005, we identified a cluster of 99 voxels in the medial part of medial pulvinar exhibiting 2 peaks (figures 2 and 3C, Table S1.3). The anterior peak activity was identified as possibly originating from the posterior choroid artery of the third ventricle whereas the posterior peak was identified as a medial pulvinar activity.

In the face>fixation contrast, at p<0.05 FWE corrected threshold, activations were located in the superior temporal gyrus, supramarginal area, Rolandic operculum bilaterally and in the left postcentral gyrus as well as left insula (figure 2, Table S1.3). At uncorrected p<0.005 threshold, a cluster of 56 voxels was observed in the third ventricle and was considered as activity from posterior choroid artery of the third ventricle (figure 2, bottom). A smaller cluster of 21 voxels was also identified in medial pulvinar. In the hand>fixation contrast, at a p<0.05 FWE corrected threshold, we identified activity in the left angular gyrus and right Rolandic operculum. At uncorrected p<0.005 threshold, we identified one cluster in the third ventricle (originating from posterior choroid artery) and one cluster in the medial pulvinar (figures 2 and 3C, Table S1.3). In the foot>fixation contrast, at p<0.05 FWE corrected threshold, activations were observed in the left superior temporal gyrus, the Rolandic operculum and the left angular gyrus (figures 2 and 3C, Table S1.3). At uncorrected p<0.005 threshold, we identified a cluster of 181 voxels. This cluster encompassed both voxels from the part of the medial pulvinar as well as voxels within the third ventricle. It exhibited 3 peaks. Two peaks were identified as bilateral activity from posterior choroid artery of the third ventricle (Li et al., 2018; Neau and Bogousslavsky, 1996) and one peak was identified as medial pulvinar. Additionally, at this same threshold (uncorrected p<0.005), we identified activations in the caudate nucleus for all contrasts.

Thus, overall, the Tactile localizer robustly activated a cortical-subcortical network involved in tactile processing (Chen et al., 2008; Grahn et al., 2008; Huang et al., 2012; Neau and Bogousslavsky, 1996).

### The pulvinar is activated by visual face and non-face stimuli and tactile hand and foot stimuli but not by auditory voice and non-voice stimuli

The goal of our study was to investigate the sensory properties of the pulvinar during passive sensory stimulation. Figure 3 summarizes the observed activations. The precise segmentation of the pulvinar was achieved thanks to the human Morel atlas (Morel et al., 1997). Visual activations in the All>Fixation contrast were widespread and encompassed the Inferior pulvinar, the ventral part of the lateral pulvinar as well as the ventral part of the medial pulvinar (p<0.05, FWE correction for multiple comparison at voxel level, figure 3B, left panel). The activation also included the MGN. These activations were bilateral. The specific face>object+scene activated a cluster of 34 voxels located in the right medial pulvinar, in its ventral anterior portion (p<0.05, FWE correction for multiple comparison at voxel level, figure 3B, right panel).

Tactile activations in the All>Fixation contrast were focal and activated a small cluster of 19 voxels in the posterior part of the left medial pulvinar (uncorrected p<0.005, figure 3C, top panel). The specific face>fixation contrast activated a cluster of 21 voxels located in this posterior part of the left medial pulvinar (Table S1.3). The specific hand>fixation contrast activated a cluster of 97 voxels located in this posterior part of the left medial pulvinar (Figure 3C, bottom left panel, Table S1.3). The specific foot>fixation contrast activated a cluster of 160 voxels located in this posterior part of the left medial pulvinar (Figure 3C, bottom right panel, Table S1.3) and a much smaller cluster of 4 voxels in the posterior part of the right medial pulvinar (Table S1.3). Two peaks could be identified in the hand and foot left activation clusters. These peaks were at identical location for both hand and foot stimulations. The peak of the face activation was located in between the two hand/foot peaks.

In order to further investigate the medial pulvinar responses to tactile stimulation, we quantified the percent signal change (PSC) in a spherical seed of 4mm of radius created from the activated cluster extracted from the All>fixation contrast analysis (MNI coordinates, x=-7, y=-30, z=5, figure 4, right panel). The goal was to compare PSCs across stimulation conditions. We tested six lateralized tactile stimulation conditions: left face, right face, left hand, right hand, left foot and right foot. Figure 4 shows a significant PSC relative to baseline for left hand stimulations (0.0971%, p<0.01, Wilcoxon signed rank test), left foot stimulations (0.0787%, p<0.01), and right face stimulations (0.0975%, p<0.008) conditions (Figure 4). Thus, the seed located in the left medial pulvinar showed significant activation by ipsilateral left hand and foot and contralateral right face stimulations. However, no significant difference was found between the six conditions (Friedman test, p=0.16, X=27.8857).

**Figure 4:**
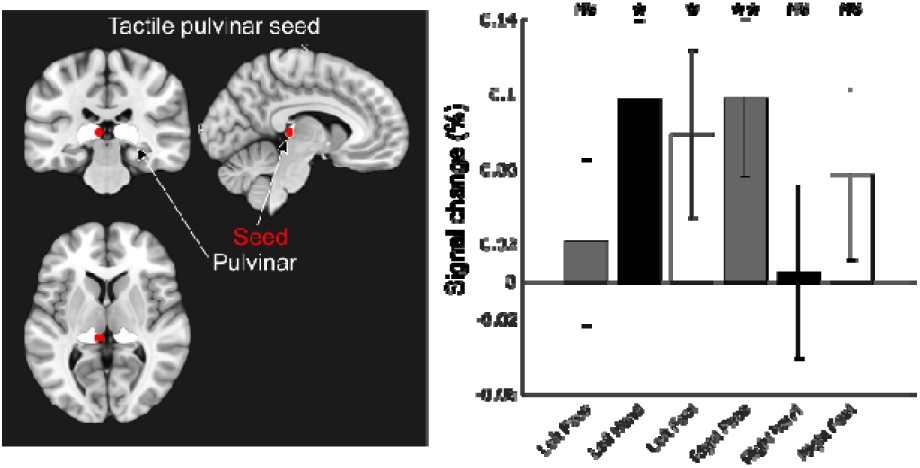
PSC of left medial pulvinar for tactile localizer conditions. Right panel. Location of the 4mm radius spherical seed in left medial pulvinar (Seed, red, MNI coordinates, x=-7, y=-30, z=5; pulvinar, white). Left panel. Percent signal change for left, right face, hand and foot tactile stimulation conditions (Wilcoxon signed rank test, *: p-value<0.05, **: p-value<0.01, corrected for multiple comparisons-Dunnett’s test).

Overall, it is thus unclear whether the pulvinar holds a tactile body map and how ipsilateral and contralateral stimulations are processed. This will have to be explored with higher resolution methods such as fast ultrasound imaging, intracortical recordings or high-field 7T imaging.

### Right ventral medial pulvinar specifically connects to the contralateral temporo-frontal ventral visual pathway during face processing

In order to assess the intrinsic connectivity affiliation of pulvinar with the rest of the brain during the face localizer task, we employed seed-based connectivity analyses using previously identified activation clusters extracted from the face localizer task as seeds. For the all>fixation contrast, two medial pulvinar clusters (one left and one right) were used as seeds and for the face>object+scene contrast, the right medial pulvinar cluster was used as seed.

Overall, in the all>fixation contrast (figure 5, left panel, tops rows, figure S1), the left seed was mostly connected to the lateral and medial prefrontal cortex and disconnected from medial posterior parietal cortex. The right seed was likewise connected to the lateral and medial prefrontal cortex. It was additionally connected to the temporal cortex and the medial part of the central sulcus. Prefrontal and central sulcus activations were grossly bilateral.

**Figure 5:**
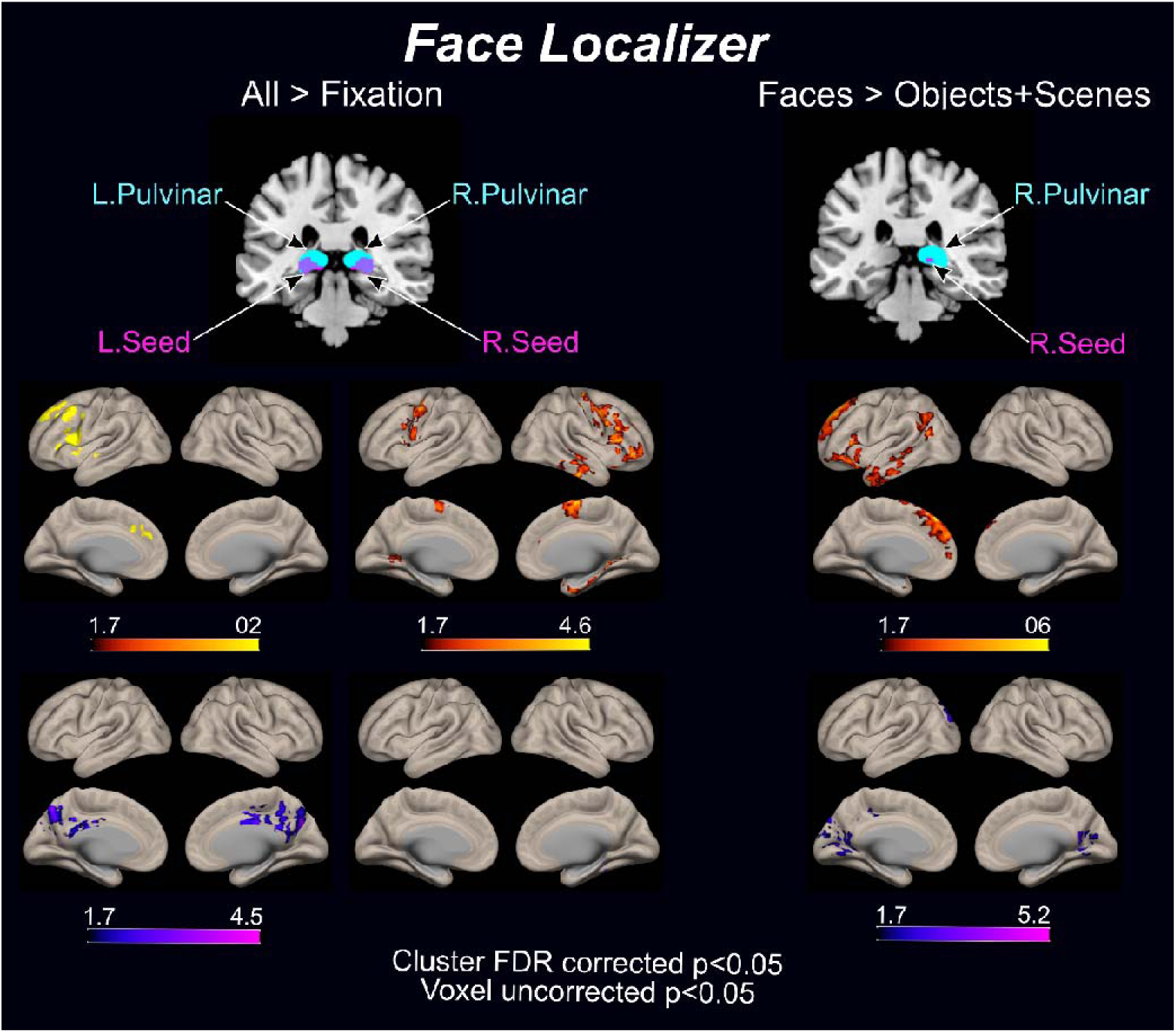
Seed-based connectivity of the pulvinar with the cortex in face localizer tasks. Top panel. Pulvinar nucleus (cyan), pulvinar seed locations in purple. Bottom panel. Seed-to-voxel connectivity analysis displayed on inflated brain representations for all>fixation contrast (left, left seed and right seed) and face>object+scene contrast (right, right seed). Maps thresholded at cluster level FDR correction p<0.05 and voxel level uncorrected p<0.05.

More specifically, the left seed was positively correlated with left frontal gyrus, left frontal pole and left precentral gyrus (figure 5, left panel, left columns, top rows, Table S2.1). It was also negatively correlated with precuneus cortex and cingulate gyrus (figure 5, left panel, left columns, bottom rows, Table S2.1). The right seed was positively correlated with right precentral gyrus, frontal gyrus, frontal pole, supplementary motor cortex, temporal gyrus, parahippocampal gyrus, temporal pole, hippocampus, orbitofrontal cortex, thalamus, and temporal gyrus as well as to left lingual gyrus (figure 5, left panel, right columns, top rows, Table S2.2).

The face>object+scene contrast mostly highlighted an increase in the right seed connectivity with the left prefrontal, occipitotemporal and temporal areas and a slight decrease in its connectivity with the medial occipital cortex bilaterally. More specifically, this right medial pulvinar seed was mainly positively correlated with left frontal pole, left frontal gyrus, left orbitofrontal cortex, left occipital cortex, left angular gyrus, left paracingulate gyrus, left temporal gyrus and left temporal pole (figure 5, right panel, top rows, Table S2.3). It was negatively correlated with different regions located mainly in the left hemisphere including the precuneus cortex, intracalcarine cortex, lingual gyrus, occipital cortex, in addition to right intracalcarine cortex (figure 5, right panel, bottom rows, Table S2.3). Thus, the seed was contralaterally connected to prefrontal, temporal and higher occipital visual areas, highlighting the ventral visual stream and it was disconnected from bilateral or contralateral lower-level occipital visual areas.

### Connectivity of the left medial medial pulvinar shifts dynamically as a function of the location of the tactile stimulation on the body

We proceeded in the same way to assess the intrinsic connectivity affiliation of pulvinar with the rest of the brain during the tactile localizer task. We thus employed seed-based connectivity analysis using the previously identified left medial pulvinar cluster activation extracted from the tactile localizer all>fixation contrast as seed. In the all>fixation contrast, there was no notable connectivity between the seed and the rest of the brain (figure 6). This could be due to an actual absence of functional connectivity between the seed and the cortex. This is unlikely. Alternatively, this could be due to the fact that each tactile stimulation activated a different cortical network. The analysis of the seed to whole brain connectivity for each of the face>fixation, hand>fixation and feet>fixation independently confirmed this latter hypothesis (figure 6, cluster level FDR correction p<0.05 and voxel level uncorrected p<0.05). Indeed, in the face contrast, the seed showed enhanced contralateral connectivity with occipital regions encoding the periphery of the visual field, and inferior parietal cortex in addition to the precuneus. In the hand contrast, the seed showed enhanced bilateral connectivity with the posterior cingulate cortex and the precuneus. In the foot contrast, the seed showed enhanced ipsilateral connectivity with motor regions in the precentral and postcentral gyrus, inferior frontal gyrus and parietal cortex. The specific foot>face contrast described enhanced connectivity with ipsilateral prefrontal cortex (inferior frontal gyrus and middle frontal gyrus) and anterior cingulate cortex in addition to the precentral gyrus and supplementary motor cortex. The specific foot>hand contrast described enhanced connectivity particularly with inferior frontal gyrus and decreased bilateral connectivity with the paracingulate cortex gyrus, anterior cingulate cortex and left inferior temporal gyrus.

**Figure 5:**
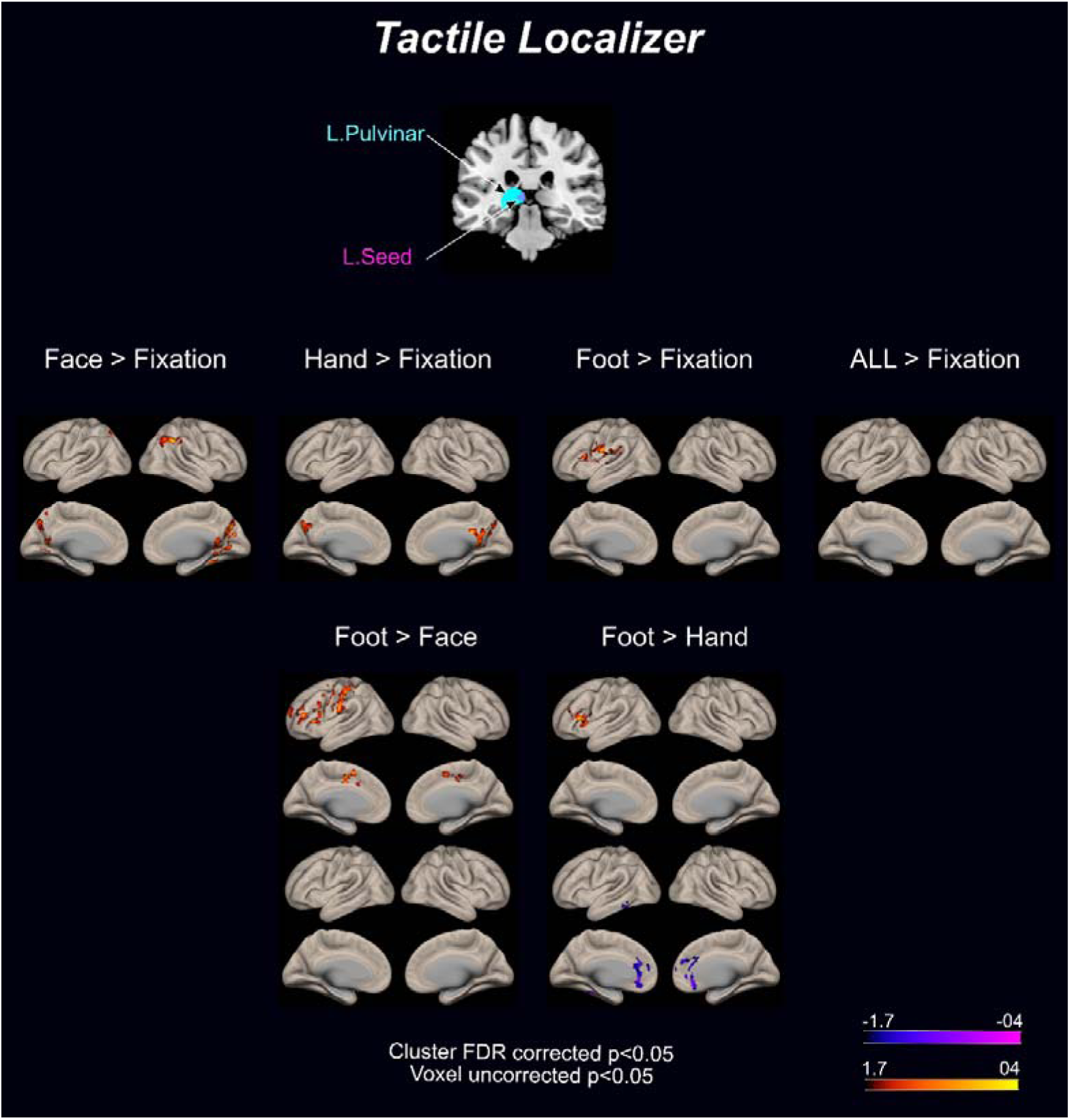
Seed-based connectivity of the pulvinar with the cortex in face localizer tasks. Top panel. Pulvinar nucleus (cyan), pulvinar seed locations in purple. Bottom panel. Seed-to-voxel connectivity analysis displayed on inflated brain representations for Face>Fixation, Hand>Fixation, Foot>Fixation, Foot>Face and Foot>Hand. Maps thresholded at cluster level FDR correction p<0.05 and voxel level uncorrected p<0.05.

## Discussion

The present study investigated the involvement of the pulvinar in sensory processing. Stimulating independently three different sensory modalities (i.e. visual, auditory and tactile) thanks to three distinct localizer tasks and using a voxel resolution (1.6×1.6×2.5mm) two or three time higher than standard acquisitions (e.g. 2.5×2.5×2.5mm or 3×3×3mm) with a field of view covering the entire brain, we confirm the pulvinar’s significant role in visual processing. We additionally demonstrated its involvement in tactile processing associated with very localized activations. Unexpectedly, we showed no significant role of the pulvinar in auditory processing. Interrogating the functional connectivity between the identified visual and tactile pulvinar regions of interest and the rest of the brain we showed that, within a given localizer task (visual or tactile), the pulvinar changed its connectivity pattern with the cortex as a function of the ongoing stimulations, thus dynamically adjusting to the sensory demand. For example, during face processing, the pulvinar exhibited decreased connectivity with low level visual processing areas and increased connectivity with cortical regions specialized in face recognition, as well as frontal areas as compared to when processing non-facial stimuli.

### Medial pulvinar face activations

Our results corroborate the role of pulvinar in face processing as previously demonstrated in various studies (Arcaro et al., 2018; Benarroch, 2015; Froesel et al., 2024a; Kaas and Lyon, 2007; Wen et al., 2023). In the cortical domain, face processing induced a large activation in occipital and temporal cortices comprising both bilateral fusiform face area (FFA) and orbital face area (OFA). No activations were observed in pSTS area which has been described as a face patch by Deen and colleagues (Deen et al., 2020). This could be due to the specificity of pSTS for face movement and motion, and to the fact that our stimuli did not have those characteristics. The ventral posterior activations we report within the pulvinar are associated with object categorization and more specifically face processing aligns with previous findings (Arcaro et al., 2018; Schwiedrzik et al., 2015; Wen et al., 2023).

### Dynamic coupling of medial pulvinar with face-related cortical areas

Our functional connectivity results bring new insights on the dynamic connectivity of pulvinar with the cortex during sensory processing. Indeed, the pulvinar showed different connectivity patterns with the brain depending on the ongoing sensory modality and stimuli category. During face processing, the pulvinar increased its connectivity with frontal and ventral visual stream areas and decreased its connectivity with low level visual processing areas. The prefrontal cortex is crucial in complex cognitive tasks, including attention, working memory, and decision-making. All these functions are engaged. This increased connectivity likely reflects the coordination between sensory processing and cognitive evaluation required for recognizing and processing faces, given their socially and emotionally significance (Arend and Zimmer, 2011). The ventral visual stream, particularly regions like the fusiform face area (FFA), is specialized for object and face recognition. The enhanced connectivity between the pulvinar and the ventral stream indicates that this nucleus might facilitate the detailed processing of facial features, integrating sensory inputs to construct a coherent percept of the face. This supports the role of the pulvinar in selectively modulating visual information flow to optimize face perception and recognition (Bridge et al., 2016). The enhanced connectivity with the prefrontal cortex supports the possible engagement of attention, working memory, and decision-making during face recognition and identification (Fuster, 2019; Levy, 2024). The reduced connectivity with low-level visual processing areas, such as the primary visual cortex (V1), suggests a shift in the pulvinar’s role from basic visual processing to more specialized, higher-order tasks during face recognition. This decrease may indicate a gating mechanism where the pulvinar suppresses irrelevant or redundant information from early visual areas to prevent sensory overload, thus allowing for more efficient processing of complex visual stimuli like faces. This selective connectivity is essential for focusing cognitive resources on the most relevant aspects of the visual input. This connectivity dynamics is expected to enhance the role of pulvinar in modulating visual information flow and coordinating between sensory and cognitive regions during face recognition (Saalmann and Kastner, 2011). The pulvinar’s dynamic connectivity changes reflect its function as a hub that prioritizes and integrates relevant sensory information while filtering out less pertinent data, facilitating efficient and accurate face recognition (Shipp, 2003).

### Auditory activations in the MGN and hippocampus but not in the pulvinar

During the voice localizer task, a large cortical activation along bilateral superior temporal gyrus and sulcus was observed, coinciding with the three main voice patches (Bodin and Belin, 2020). No pulvinar activation was observed while subcortical rMGN and hippocampus were activated for both voice and non-voice conditions. This lack of activation in the pulvinar is surprising knowing that the pulvinar has been shown to be a part of the auditory system (Kaas and Hackett, 2000) and is activated by vocalizations in marmoset (Jafari et al., 2023) and macaques (Froesel et al., 2024a). The absence of pulvinar activation might be attributed to the passive nature of the task, not requiring any specific cognitive engagement from the subjects except for fixation. Alternatively, this could be due to the fact that the sounds were not delivered in a multisensory context. This point will have to be explored in future experiments involving tasks requiring active auditory discrimination. Indeed, it has been proposed that the pulvinar is more likely to be recruited during more challenging tasks in order to enhance stimuli perception (Froesel et al., 2021).

The rMGN is part of the auditory thalamus and plays a crucial role in the relay and processing of auditory information from the ear to the auditory cortex. Its activation in both voice and non-voice conditions is consistent with its well-established function in basic auditory signal transmission and initial processing (Lee and Sherman, 2011; Winer, 2006). Activation of bilateral hippocampus for both face and voice localizer tasks is consistent with previous literature. Sliwa et al, 2016 (Sliwa et al., 2016), identified neurons responding for social visual and auditory stimuli in macaque hippocampus. The identified cluster at lower statistical threshold for voice localizer than face localizer is also consistent with their findings of predominance for visual stimuli. The activation peaks we reported were close to hippocampus/amygdala border, fitting with an fMRI visual localizer study that reports interindividual variability in the localization of visual activations in the amygdala or the hippocampus (Schwarz et al., 2019). The hippocampus is known for its role in forming and retrieving memories (Suzuki and Eichenbaum, 2000). The activation of the hippocampus in response to both voice and non-voice stimuli possibly indicates the engagement of memory processes for the recognition and contextualization of auditory stimuli, regardless of their nature (Poremba et al., 2004; Sliwa et al., 2016).

### Medial pulvinar activations are similar for face, hand and foot tactile stimulations

The tactile localizer robustly activated cortical regions involved in somatosensory processing. We report activations in parietal and temporal areas, as well as bilateral activations of secondary somatosensory cortex (SII) in agreement with previous studies (Chen et al., 2008; Fujiwara et al., 2002; Hoechstetter et al., 2000). We did not observe any activation in SI. This is usually the case in studies requiring some level of attention to the tactile stimulation (Chen et al., 2008; Fujiwara et al., 2002; Hoechstetter et al., 2000), and matches our instruction to the subjects to focus on the tactile stimulations as they were presented.

The caudate nucleus of the striatum was activated across all conditions in tactile localizer at low statistical threshold. This nucleus is one of the main input to basal ganglia and receives connections from a large part of the cortex including somatosensory and motor cortex (Grahn et al., 2008). Functionally, it is implicated in high level cognitive process like goal planning, sensorimotor processes and oculomotor circuit (Grahn et al., 2008). During the tactile localizer task, participants had to focus on the stimulated body part and press a response button when fixation point was changing color. Thus, activation of the caudate nucleus in this task might be due to either its anatomical connections with somatosensory cortex or the oculomotor demands or a combination of both.

Our first whole brain contrast analysis showed a weak activation cluster in the medial part of medial pulvinar during tactile processing of air puffs. The PSC analysis confirmed the increase in activity in medial pulvinar during face, hand, and foot stimulations and showed no difference between the three conditions. This result is, to our knowledge, the first demonstration of pulvinar activation following puff stimulation in humans and reveals no body-part specific organization within the pulvinar. These results are in line with a recent fMRI study (Charyasz et al., 2023) using both passive and active tactile tasks to investigate the sensorimotor activations in the thalamus. The authors found activation in PuA, PuM and PuL in both of these tasks involving either movement or stimulation of the right finger. We add to this study PuM activations following the stimulation of other body parts. They observed pulvinar single subject activations in a restricted number of subjects for the 4 subdivisions of pulvinar but with dominance of the left over the right pulvinar. A convergent observation between the two projects is that both observed activations in left medial pulvinar during tactile stimulation. However, Charyasz et al. observed only few voxels activated in medial, lateral and inferior pulvinar and a high number of voxels in anterior pulvinar whereas we only observed activations in medial pulvinar. The location of the observed voxels within medial pulvinar across the two studies is also different, our activations being located close to the medial border of the pulvinar whereas they observed voxels more laterally and anteriorly. This might be due to the difference in the type of tactile stimulation applied, one study using air puffs and the other study using air pulses through an inflatable finger clip.

The identified pulvinar tactile activation is located within the pulvinar and the peak of the cluster falls within the Morel pulvinar ROI (Morel et al., 1997). This is intriguing as no direct anatomical connections exist between the pulvinar and the somatosensory cortex. Three hypotheses may account for this activity. First, the activity might be miss located in the pulvinar while originating from a nearby structure. Two main candidates would be ventral posterior lateral (VPL) and central lateral (CL) nuclei of thalamus (Delhaye et al., 2018; Kumar et al., 2017; Mai and Majtanik, 2019). The VPL which relays somatosensory information to the cortex, can be dismissed due to its distance from the tactile cluster location (Delhaye et al., 2018). The CL part of the internal medullary lamina complex (Kumar et al., 2017) is closer to our tactile cluster than VPL. However, the spatial resolution of our fMRI scans allows us to unambiguously describe no overlap between CL and the reported tactile cluster. Additionally the CL is involved in pain perception (Nowacki et al., 2023) and air-puff stimulation used in the tactile localizer did not elicit any pain perception in none of the subjects. Thus, we can rule out the hypothesis of adjacent structures accounting for the activity observed in the posterior ventral cluster. Another possibility is that projections between our pulvinar cluster and somatosensory system may exist but have not yet been described. The cluster is located close to the third ventricle and injections and recordings are difficult to target at that location. Yet, the most plausible hypothesis is that of an indirect activation of medial pulvinar through secondary projections possibly related to awareness of tactile perception. Pulvinar is indeed implicated in visual attention and saliency. Participants were instructed to focus on the tactile stimulations. Thus, the pulvinar may have been activated due to the deployment of attentional processes or to the intrinsic salience of the unpredictable tactile stimuli (Froesel et al., 2021; Guedj and Vuilleumier, 2020). This possibly suggests that the pulvinar responds to general sensory salience rather than to the specific nature of the stimulus. This aligns with the pulvinar’s role in modulating attention towards stimuli that require immediate processing, regardless of their specific sensory characteristics highlighting its function in integrating diverse sensory information and directing attention towards salient stimuli, irrespective of their specific nature. Interestingly, we found activation in the medial part of the medial pulvinar and not in anterior pulvinar (PuA) in spite of the fact that anatomical connections between PuA and lateral and anterior parts of the parietal cortex and area 1 have previously been described (Delhaye et al., 2018). This might be due to the fact that our localizer did not activate these cortical regions.

### Dynamic coupling of the tactile medial pulvinar ROI with cortical areas

Connectivity analysis showed that the tactile medial pulvinar seed cluster exhibited distinct connectivity patterns depending on the location of the tactile stimulation. Indeed, while in the all>fixation contrast, no significant connectivity was observed, the separate contrasts (face>Fixation, Hands>Fixation and feet>Fixation) resulted in distinct non-overlapping cortical functional connectivity patterns with this tactile medial pulvinar ROI. This indicates that tactile stimulation resulted in different pulvino-cortical connectivity patterns as a function of which body part was stimulated.

Within the face>Fixation contrast, the left medial pulvinar exhibited enhanced contralateral connectivity with occipital regions involved in visual field periphery processing (Tootell et al., 1998), the inferior parietal cortex, and the precuneus. These findings align with the pulvinar’s role in integrating multisensory information, particularly in coordinating visual and tactile inputs (Benarroch, 2015; Fiebelkorn et al., 2019; Saalmann and Kastner, 2011), as well as with privileged convergence between tactile information and visual peripheral information (Guipponi et al., 2015). The connection with the inferior parietal cortex, a region associated with attention and sensorimotor integration (Behrmann et al., 2004; Sereno and Huang, 2014), suggests that the pulvinar may facilitate the alignment of visual and somatosensory maps, supporting efficient cross-modal processing.

The same seed showed for Hands>Fixation contrast, enhanced bilateral connectivity with the posterior cingulate cortex and the precuneus. The posterior cingulate cortex is a key node in the default mode network (DMN), which is implicated in self-referential thought and internally focused attention (Raichle, 2015). The precuneus, another DMN region, is involved in visuospatial imagery and attention (Raichle, 2015). The pulvinar’s connectivity with these regions during hand stimulation may reflect the integration of tactile information with internally directed cognitive processes, potentially facilitating the mental representation of hand movements or positions.

In the Feet>Fixation contrast, this left medial cluster demonstrated an enhanced ipsilateral connectivity with motor regions (precentral and postcentral gyri), the inferior frontal gyrus, and the parietal cortex. The involvement of motor regions suggests that the pulvinar plays a role in linking tactile inputs from the feet with motor planning and execution areas, which may be crucial for tasks requiring coordination between sensory input and motor output, such as walking or balance. The connectivity with the inferior frontal gyrus, a region associated with motor planning (Dick et al., 2019; Niedermeyer, 1998), further supports the pulvinar’s role in preparing and executing motor responses based on tactile information.

The specific contrasts between foot and face, as well as foot and hand, revealed additional insights into the functional specialization of the pulvinar’s connectivity. The foot>face contrast showed enhanced connectivity with ipsilateral prefrontal regions (inferior and middle frontal gyri), the anterior cingulate cortex, and motor areas (precentral gyrus and supplementary motor area). These connections highlight the pulvinar’s involvement in integrating tactile information from the feet with higher-order executive functions, such as decision-making and motor planning. The anterior cingulate cortex’s involvement suggests a role in monitoring and adjusting behavior in response to tactile stimuli from the feet, which is critical for maintaining balance and posture. In the foot>hand contrast, there was enhanced connectivity particularly with the inferior frontal gyrus and decreased bilateral connectivity with the paracingulate gyrus, anterior cingulate cortex, and left inferior temporal gyrus. This pattern indicates a shift in the pulvinar’s role from integrating tactile information with motor planning (in the case of foot stimulation) to modulating higher cognitive functions, such as attention and decision-making, when the tactile input comes from the hands. The decreased connectivity with regions involved in self-referential processing and visual memory (paracingulate gyrus and inferior temporal gyrus) suggests that the pulvinar’s connectivity is finely tuned to the specific demands of the tactile task at hand.

### Relation with previous work

Guedj and Vuilleumier proposed five clusters functional parcellation of the pulvinar based on fMRI resting-state data from 100 healthy subjects (Guedj and Vuilleumier, 2020). According to their parcellation, our face localizer activation cluster is located in their dorsomedial cluster. Our tactile localizer activation cluster is located in their ventromedial cluster. This supports the hypothesis of a process dependent organization within pulvinar. Guedj and Vuilleumier further specify resting-state connectivity of each cluster with the rest of the brain as well as task-related connectivity. The dorsomedial cluster was identified to be connected with cortical regions involved in default mode and saliency networks. The ventromedial cluster showed a widespread connectivity pattern including the ventral visual extrastriate areas and medial temporal lobe which belong to the visual ventral stream important for object recognition including face recognition. Our activations are thus not consistent with the connectivity of the corresponding cluster described by Guedj and Vuilleumier.

Taken together, these results advance our understanding of the pulvinar’s multisensory role in the brain, highlighting its role in face processing and tactile processing possibly supporting its role in integrating and modulating sensory information to support higher-order cognitive functions and attentional processes. Future studies should address the functional parcellation of pulvinar and their contribution to different cognitive functions so as to clarify the role of this nucleus in human cognition.

## Method

### Participants and ethical approval

Twenty-five subjects aged between 20 to 33 years old (12 males and 13 females) with mean age of 28.08 years old (SD = 3.8) participated in the study after signing an informed written consent. All subjects were healthy and had no history of psychiatric or neurological disorders and had normal or corrected-to-normal vision. Edinburgh Handedness Inventory laterality test (doi: https://doi.org/10.1016%2F0028-3932%2871%2990067-4) is used to determine a lateralization score for each participant (17 right-handed, 1 left-handed and 7 ambidextrous). The present study was approved by the French ethical comity CPP sud-Med IV (Comité de Protection des Personnes Sud-Méditerranée IV - 19.00758.190401-MS02 – 2018-A03438-47).

### Experimental setup

During MRI sessions, participants were lying on their back in a supine position. A mirror was positioned above their head and fixated to the head coil. A plastic translucent screen of 30 cm of radius was set at 1 meter from the eyes/mirrors. Visual stimuli were retro-projected onto this screen. Tactile stimulation was delivered by air pressure at 2.3 bars (air puffs) through plastic tubes rigidly maintained close to the target body locations. This level of pressure induced in all participants a clear and unambiguous perception of tactile stimulation on the target area. Air pressure was created by a compressor (*SIP Industrial*, SIP QT 50/10 Low Noise Direct Drive Compressor) and adjusted by pressure regulators placed outside of the scanner room. Auditory stimuli were delivered using an MRI compatible device (Sensimetric Store, Model S14 - Insert Earphones for fMRI Research). Volume adjustment was checked during installation of the participant. Eye position (x and y for right or left eye) was recorded using an eye tracking system (Eyelink 1000, SR Research™) and sampled at 1000Hz. Stimulus control for all modalities as well as online eye position control was achieved using Presentation software (Neurobehavioral systems™) coupled with and I/O control Ardiuno-based setup.

### Behavioral tasks

Subjects performed three different localizer tasks. In all three localizer tasks, in order to have participants keep focusing on the stimuli during the entire runs, they were asked to fixate the central point displayed on top of the stimuli and to answer by pressing a button when the point change color.

#### Face localizer task (**Fig. 1**, top panel)

The face localizer task was composed of four conditions: fixation, human faces, objects (cars) and scenes (corridors). Stimuli dataset is based on the open fLoc (http://vpnl.stanford.edu/fLoc/) functional localizer package (Stigliani et al., 2015). During the task, subjects were asked to maintain their gaze on a fixation point presented at the center of the screen, overlaying the presented images. Subjects had to perform four runs of the face localizer. Each run was composed of a total of 24 blocks organized in 6 sets of the 4 conditions or blocks of stimuli (Fixation, Faces, Objects, Scenes). Conditions were pseudo-randomized within each set. Each block was composed of 16 different consecutive images each displayed for 500ms and selected randomly from a set of 144 images. Images were presented in the center of the screen and were 24° x 24°. In each run, the task started after 5 pulses to insure stability of magnetic field.

#### Tactile localizer task (**Fig 1**. Middle panel)

The tactile localizer task is composed of seven conditions: fixation, tactile stimulation to the left cheek, to the right cheek, to the left hand, to the right hand, to the left and right feet. During this task, participants were asked to perform four runs each composed by a total of 35 blocks organized in 5 sets or repetitions of the 7 conditions pseudo-randomized within each set. One block was composed by 5 consecutives tactile stimulations (air puffs) of 300ms duration with a random inter-stimulation interval between 1000 and 2000ms. Tactile stimulation was not associated with any simultaneous auditory stimuli as compressor and regulators were placed outside of the scanner room. Targeted locations were maintained as similar as possible across subjects: below the cheekbone for cheeks stimulation, half way between the ear and the nose; between index and thumb for hand stimulation and on the top of the foot for feet stimulation, aligned with the shin, and half way between the ankle and the fingers.

#### Voice localizer task (**Fig. 1**, bottom row)

The voice localizer task is composed of three conditions: fixation, voices and non-voice sounds. Stimuli dataset (one dataset of voice, one dataset of non-voice, 20 audio sequences per set) was taken from (Pernet et al., 2015) and (Belin et al., 2000). For the description of the stimuli in the blocks see Table 1 of Pernet et al., 2015. Participant were asked to perform one run of 72 blocks lasting 8 seconds. The three conditions were played equally (24 voice blocks, 24 non-voice blocks, 24 fixation blocks) and were presented randomly during the run.

### Image acquisition

MRI data were acquired with a Siemens 3T PRISMA scanner and using a circularly polarized head coil of 64-channels (Siemens Healthineers, Erlangen, Germany). High-resolution anatomical images were acquired: T1-weighted images MPRAGE sequence for the T1-weighted (sagittal incidence, GRAPPA factor=2, FOV=180×224X256 mm, 0.8 mm isotropic voxel size, TR=3.0 s, TE=3.7ms, TI=1.1s, flip angle =8 LJ). High-resolution 3D-T2-weighted anatomical images were acquired with SPACE sequence (sagittal incidence, GRAPPA factor=3, FOV=180×224X256 mm, 0.8 mm isotropic voxel size, TR=3.0s, TEeff=299 ms). Functional images were acquired with EPI sequence covering whole cortex (50 axial slices, multiband factor=2, FOV=204×204×125, voxel size=1.6×1.6×2.5 mm with phase encoding along AP direction, TR = 2.3s; TE = 36 ms; FA = 80°). To correct for geometrical distortion, a magnetic field map was acquired before the functional runs with Siemens gre_field_mapping sequence (44 axial slices, FOV=225×225×132, voxel size=2.5×2.5×3.0 mm TR =500ms, TEs = 4.59 and 7.05ms, FA = 60 LJ). The face localizer task involves 4 runs of 100 volumes each, the tactile localizer task involves 4 runs 120 volumes and the voice localizer task involved 1 run of 264 volumes.

### Image preprocessing

Image pre-processing was performed using Statistical Parametric Mapping (SPM12) (Friston et al., 1994) software (Wellcome Centre for Human Neuroimaging, University College London, UK; http://www.fil.ion.ucl.ac.uk/spm). First, the first six volumes of the task were discarded allowing for T1 equilibrium. To adjust within-volume time differences, temporal misalignment between different slices of the functional data, introduced by the sequential nature of the fMRI acquisition protocol, is corrected using SPM12 slice-timing correction (STC) procedure (Henson et al. 1999), where the functional data is time-shifted and resampled using sinc-interpolation to match the time in the middle of each TA (acquisition time). Functional data were then realigned using SPM12 realign & unwarp procedure (Andersson et al. 2001), where all scans are resampled to a reference image (first scan of the first session) using b-spline interpolation. This procedure also addresses potential susceptibility distortion-by-motion interactions by estimating the derivatives of the deformation field with respect to head movement and resampling the functional data to match the deformation field of the reference image. Finally, functional scans were co-registered with the 3D structural image and spatially normalized against the standard, stereotactic space of the MNI template. Spatial smoothing was performed with a 6-mm full with half maximum Gaussian kernel.

### Functional MRI GLM analyses

Functional images were analyzed using statistical parametric mapping software (SPM12) (Friston et al., 1994, Wellcome Centre for Human Neuroimaging, University College London, UK; http://www.fil.ion.ucl.ac.uk/spm). The first five functional volumes were discarded to allow for T1 equilibrium. At single subject level, a fixed-effect model was created for each individual participant in order to perform random-effect analyses with FWE p<0.05 corrected for multiple comparisons. The motion parameters were included in the General Linear Model (GLM) as regressors to correct the head movement. In order to assess brain activations specific to the experimental conditions, different contrasts were computed for each task. For the face localizer task, activations related to the 3 visual conditions were assessed by all>fixation contrasts. Activations specifically related to the face condition was assessed by the following contrasts: face>objects+scenes. For the voice localizer task, non-specific, voice-specific and non-voice-specific auditory activations were assessed by the following contrasts: voice>fixation, non-voice>fixation and voice>non-voice. For the tactile localizer task, activations related to tactile stimulation of face (cheek), hands, and feet were assessed by the following contrasts: face>fixation, hand>fixation, foot fixation. For the face and the tactile localizer tasks, in the GLM, contrasts were replicated and rescaled over runs accounting for inter-run variability by applying the same contrast weights to the corresponding regressors across the multiple runs and scaling them appropriately. This option allows to ensure that the contrasts are appropriately normalized and average the effects across the runs, enhancing the statistical power to detect consistent effects. A second-level random-effects analysis was then conducted and statistical results were thresholded using a voxel-wise family-wise error (FWE) corrected height threshold of p < 0.05. For tactile localizer percent signal change was calculated. A 4mm radius spherical region of interest was created using marsbar matlab toolbox ((Brett et al., n.d.)) and located at MNI coordinates of x=-7, y=-30, z=5, to as to encompass the pulvinar All>fixation tactile activation cluster identified in whole brain analysis..

### Seed Based analyses

In order to establish the intrinsic connectivity network affiliation of the pulvinar within each localizer (Face, Voice and Tactile), a seed to voxel analysis using the CONN19b toolbox (Whitfield-Gabrieli and Nieto-Castanon, 2012) was used. First, we extracted pulvinar activation clusters by applying the Morel pulvinar mask at FWE p<0.05 corrected for multiple comparisons (Morel et al., 1997; Niemann et al., 2000). For face localizer task, seeds from the following contrasts were extracted: all>fixation (left seed, x=-21, y=-28, z=-2, size=232 voxels, right seed, x=21, y=-28, z=-3, size=278 voxels), face>object+scenes (x=15, y=-32, z=0, size=24 voxels). Then, in the CONN toolbox, additional denoising steps were carried out. Potential outlier scans were identified from the observed global BOLD signal as well as the amount of subject-motion in the scanner. Principal components were brought out from white matter and cerebrospinal fluid time-series to reduce noise in reference to the anatomical CompCor approach (Component-based Noise Correction) for each participant (Behzadi et al., 2007). These components, in addition to head movements parameters were used as confounds in the denoising step. The signal BOLD time-series were extracted and band pass-filtered for each participant (0.008–0.09 Hz). Finally, all pulvinar clusters extracted from SPM contrasts within each localizer task were implemented in the CONN toolbox as ROIs. These were then used unilaterally as seeds in the seed to voxel analysis during which correlation maps on voxel-level were computed by correlating the BOLD-signal time course, averaged over all voxels within identified regions with the BOLD-signal time-course of every single voxel in the brain over the duration of the task for each participant. For the group-level analysis the Conn toolbox used ReML estimation of covariance components (which are evaluated through F-statistical parameter maps) to implement contrasts for analyses at the voxel level as repeated-measures analyses (Mueller et al., 2015). We reported the correlation maps within the contrasts that have served to extract the seed clusters. A threshold at the clusters p < 0.05 FDR corrected and peak voxel threshold p < 0.05 uncorrected were used.

## Abbreviations

PuI: inferior pulvinar (green), PuA: anterior pulvinar (red), PuL: lateral pulvinar (blue), PuM: medial pulvinar (white), MD: mediodorsal nucleus (brown), CM: centre median nucleus (dark green), CL: central lateral nucleus (cyan), VPL: ventral posterior lateral nucleus (gold), MGN: medial geniculate nucleus (orange), LGN: lateral geniculate nucleus (yellow).

## Acknowledgements

S.B.H. was funded by the French National Research Agency (ANR) ANR-16-CE37-0009-01 grant and the LABEX CORTEX funding (ANR-11-LABX-0042) from the Université de Lyon, within the program Investissements d’Avenir (ANR-11-IDEX-0007) operated by the French National Research Agency (ANR). We also thank Thomas Perret, Johan Pacquit, and Marco Bimbi for help with the hardware computational resources.

## Author contributions statement

Conceptualization: S.B.H., M.F.; Data Curation: MGT.M., M.F. and S.D.; Formal Analysis: MGT.M., M.F. S.D. and S.B.H; Funding Acquisition: S.B.H.; Investigation: MGT.M., M.F., S.D. and S.B.H.; Methodology: MGT.M., M.F., S.D. and S.B.H.; Supervision: S.B.H.; Writing—Original draft: MGT.M., M.F and S.D and S.B.H.; and Writing—review & editing: MGT.M., M.F., S.D. and S.B.H.

### Additional information

Data and code are available upon request.

## Supplementary material

**Supplementary figure S1.**
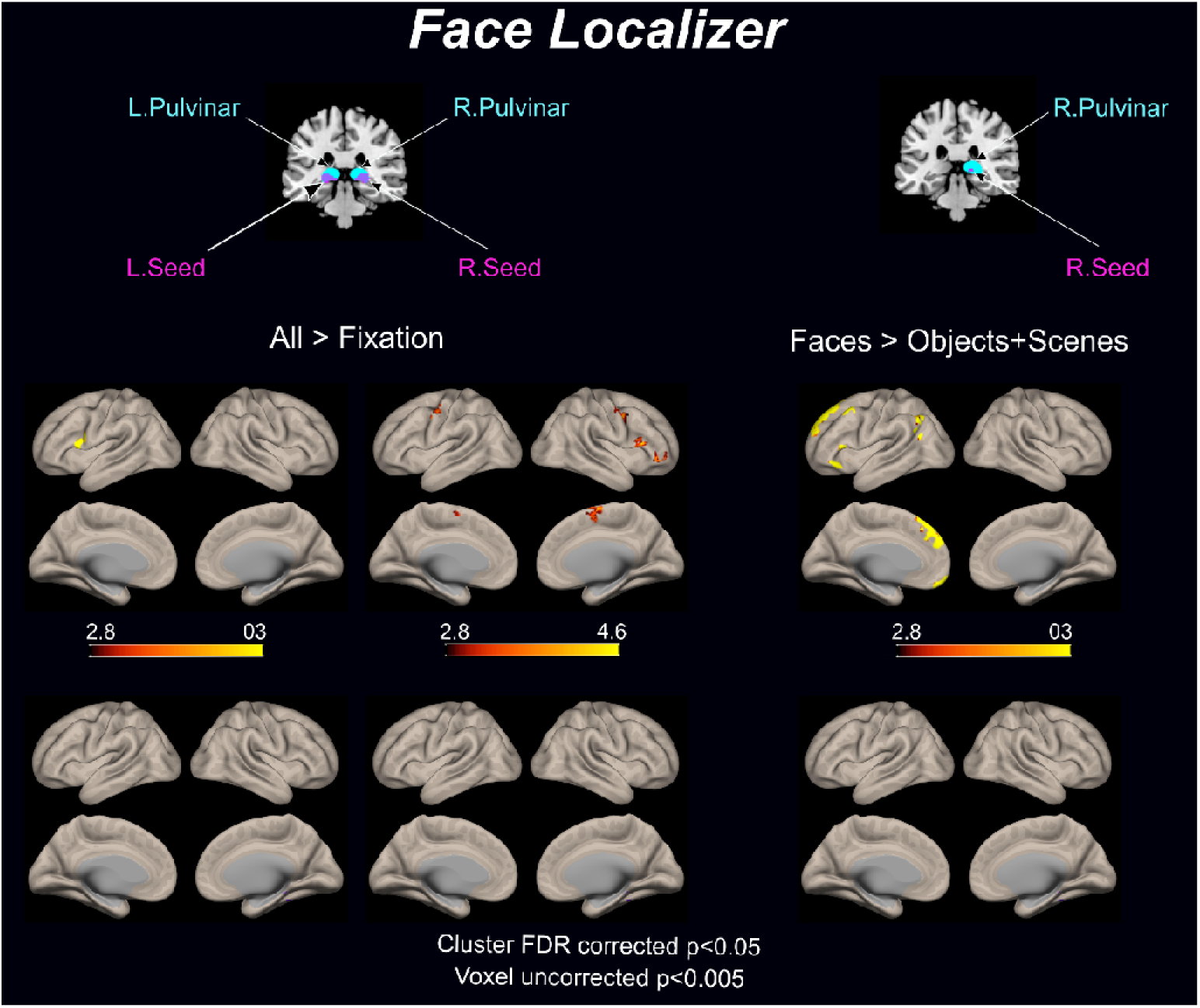
Seed-based connectivity of the pulvinar with the cortical domain (cluster FDR p<0.05 and voxel uncorrected p<0.005). Same data as in Figure 5, except threshold is more restrictive in figure S1.

**Supplementary Table S1.1.**
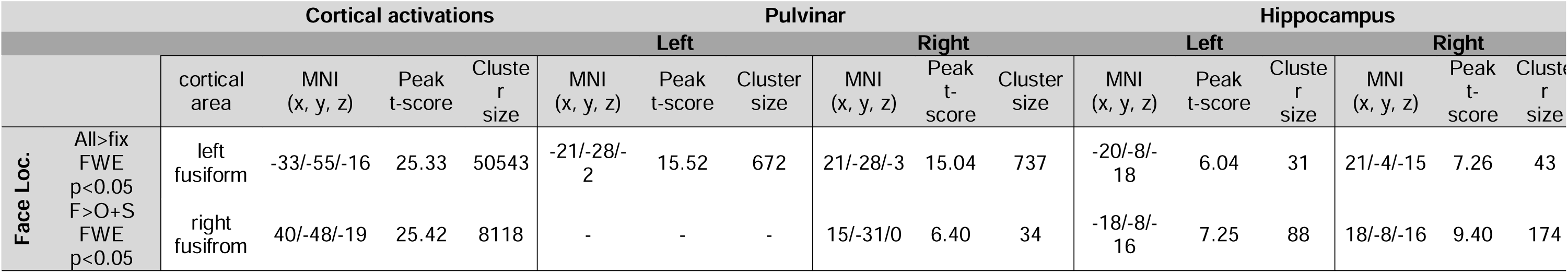
fMRI activations in face localizer task. Cortical, pulvinar and hippocampal regions displaying cluster activations in the face localizer task, for the All>Fixation contrast and the Face>Objects+Scenes contrast. Cluster peak location, peak t-score and cluster size are indicated. Star (*) indicates presence of two local peaks within the same cluster.

**Supplementary Table S1.2.**
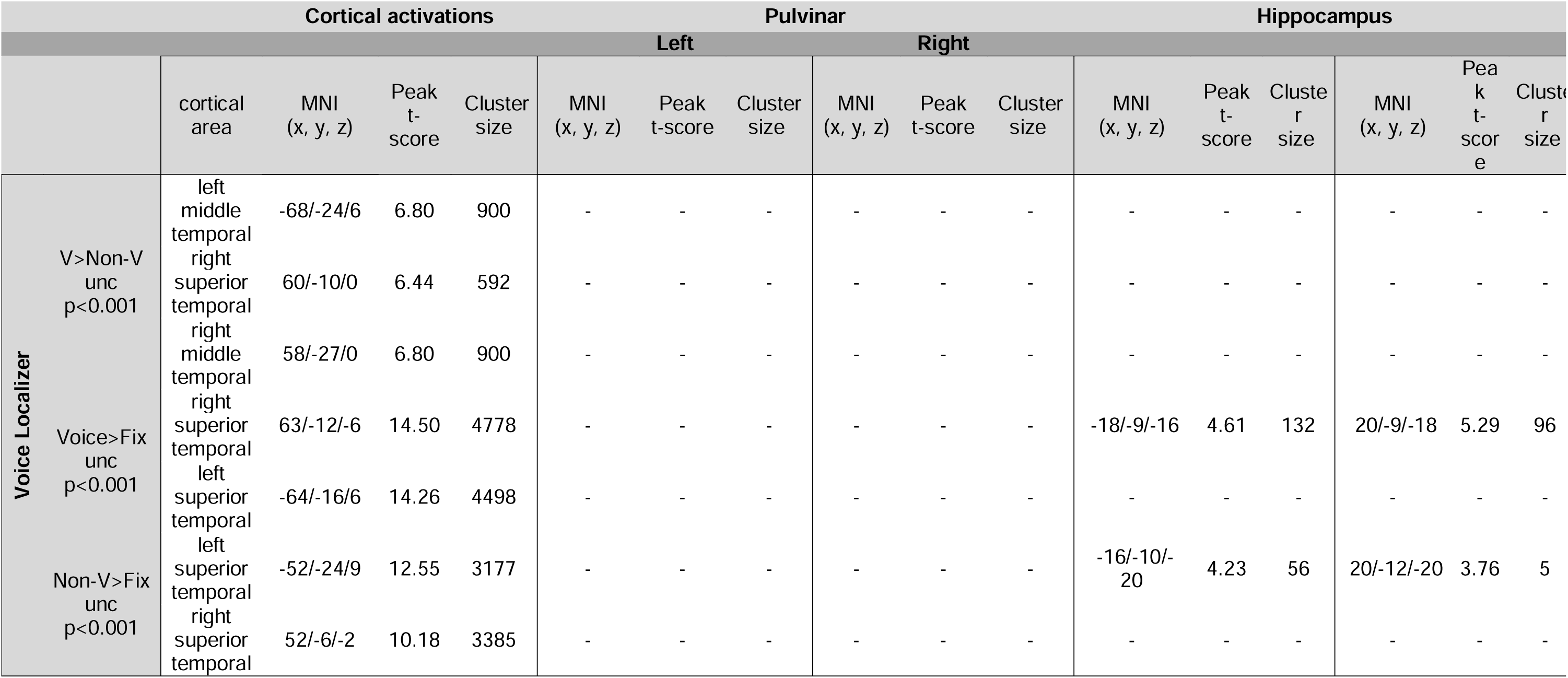
fMRI activations in voice localizer task. Cortical, pulvinar and hippocampal regions displaying cluster activations in the voice localizer task, for the Voice>non-voice contrast (V>Non-V), Voice>Fixation contrast (Voice>Fix) and Non-Voice>Fixation contrast (Non-V>Fix). Cluster peak location, peak t-score and cluster size are indicated. Star (*) indicates presence of two local peaks within the same cluster.

**Supplementary Table S1.3.** fMRI activations in tactile localizer task. Cortical, pulvinar and hippocampal regions displaying cluster activations in the tactile localizer task, for the All>Fixation contrast, Face>Fixation contrast, Hand>Fixation contrast and Foot>Fixation contrast. Cluster peak location, peak t-score and cluster size are indicated. Star (*) indicates presence of two local peaks within the same cluster.

|  |  | Cortical activations |  |  |  | Pulvinar |  |  |  |  |  | Hippocampus |  |  |  |  |  |
| --- | --- | --- | --- | --- | --- | --- | --- | --- | --- | --- | --- | --- | --- | --- | --- | --- | --- |
|  |  | cortical area | MNI (x, y, z) | Peak t-score | Cluster size | Left |  |  | Right |  |  | MNI (x, y, z) | Peak t-score | Cluster size | MNI (x, y, z) | Peak t-score | Cluster size |
|  |  |  |  |  |  | MNI (x, y, z) | Peak t-score | Cluster size | MNI (x, y, z) | Peak t-score | Cluster size |  |  |  |  |  |  |
| Tactile Localizer | All>Fix unc p<0.005 | left superior temporal | -48/-34/24 | 7.03 | 330 | -3/-27/9 | 5.21 | 19 | - | - | - | - | - | - | - | - | - |
|  |  | right rolandic operculum | 51/-28/22 | 6.99 | 492 | -6/-33/3 | 6.02 | 19 | - | - | - | - | - | - | - | - | - |
|  |  | left angular | -50/-64/33 | 5.25 | 64 | - | - | - | - | - | - | - | - | - | - | - | - |
|  |  | left postcentral | -58/-16/18 | 5.12 | 37 | - | - | - | - | - | - | - | - | - | - | - | - |
|  | Face>Fix unc p<0.005 | left superior temporal | -50/-34/22 | 7.96 | 1532 | -3/-25.5/9 | 5.33 | 21 | - | - | - | - | - | - | - | - | - |
|  |  | right superior temporal | 51/-32/20 | 7.00 | 1038 | - | - | - | - | - | - | - | - | - | - | - | - |
|  |  | left insula | -38/-3/-2 | 5.99 | 32 | - | - | - | - | - | - | - | - | - | - | - | - |
|  | Hand>Fix unc p<0.005 | left angular | -50/-64/32 | 5.53 | 36 | -3/-27/9 | 6.56 | 97* | - | - | - | - | - | - | - | - | - |
|  |  | right rolandic operculum | 52/-26/22 | 5.31 | 19 | -6/-31.5/3 | 5.9 | 97* | - | - | - | - | - | - | - | - | - |
|  | Foot>Fix unc p<0.005 | left superior temporal | -51/-33/24 | 6.27 | 145 | -3/-27/9 | 7.36 | 160* | 6/-30/4.5 | 4.80 | 4 | - | - | - | - | - | - |
|  |  | right rolandic operculum | 51/-28/24 | 5.95 | 139 | -6/-31.5/3 | 6.72 | 160* | - | - | - | - | - | - | - | - | - |
|  |  | left angular | -50/-64/33 | 5.41 | 29 | - | - | - | - | - | - | - | - | - | - | - | - |

**Supplementary Table S2.1.** Connectivity results for All>Fixation face localizer contrast, left seed. Positive and negative brain connectivity for selected seed, number of voxels and % of areas activated.

| Positive connectivity |  |  |  | Negative connectivity |  |  |
| --- | --- | --- | --- | --- | --- | --- |
| All>fixation<br>Left Seed | Brain area | # voxels | % | Brain area | # voxels | % |
|  | left-middle frontal gyrus | 540 | 18 | precuneus cortex | 1424 | 25 |
|  | left inferior frontal gyrus pars opercularis | 401 | 52 | cingulate gyrus posterior division | 481 | 20 |
|  | left frontal pole | 272 | 4 | cingulate gyrus anterior division | 102 | 4 |
|  | left precentral gyrus | 254 | 6 | - | - | - |
|  | left superior frontal gyrus | 217 | 8 | - | - | - |
|  | inferior frontal gyrus left pars triangularis | 99 | 15 | - | - | - |

**Supplementary Table S2.2.** Connectivity results for All>Fixation face localizer contrast, right seed. Positive and negative brain connectivity for selected seed, number of voxels and % of areas activated.

| Positive connectivity |  |  |  | Negative connectivity |  |  |
| --- | --- | --- | --- | --- | --- | --- |
| All>fixation<br>Right Seed | Brain area | # voxels | % | Brain area | # voxels | % |
|  | bilateral precentral gyrus | 786/921 | 18/22 | - | - | - |
|  | right middle frontal gyrus | 653 | 24 | - | - | - |
|  | right frontal pole | 425 | 5 | - | - | - |
|  | supplementary motor cortex bilateral juxtapositional lobule | 143/344 | 22/48 | - | - | - |
|  | right middle temporal gyrus posterior division | 311 | 23 | - | - | - |
|  | right inferior frontal gyrus pars opercularis | 295 | 43 | - | - | - |
|  | right parahippocampal gyrus anterior division | 207 | 32 | - | - | - |
|  | right temporal pole | 202 | 8 | - | - | - |
|  | right hippocampus | 153 | 22 | - | - | - |
|  | right orbitofrontal cortex | 151 | 10 | - | - | - |
|  | right superior frontal gyrus | 134 | 5 | - | - | - |
|  | right thalamus | 116 | 9 | - | - | - |
|  | right inferior frontal gyrus pars triangularis | 106 | 19 | - | - | - |
|  | left lingual gyrus | 103 | 7 | - | - | - |
|  | right inferior temporal gyrus posterior division | 101 | 11 | - | - | - |

**Supplementary Table S2.3.** Connectivity results for Face>object+scene face localizer contrast, right seed. Positive and negative brain connectivity for selected seed, number of voxels and % of areas activated.

| Positive connectivity |  |  |  | Negative connectivity |  |  |
| --- | --- | --- | --- | --- | --- | --- |
| Faces>objects+scenes | Brain area | # voxels | % | Brain area | # voxels | % |
|  | left frontal pole | 1466 | 21 | precuneus Cortex | 314 | 6 |
|  | left superior frontal gyrus | 1006 | 36 | left intracalcarine Cortex | 167 | 26 |
|  | left orbitofrontal cortex | 520 | 31 | left lingual gyrus | 150 | 10 |
|  | left lateral occipital cortex superior division | 479 | 10 | left superior division lateral occipital cortex | 146 | 3 |
|  | left angular gyrus | 417 | 44 | right intracalcarine cortex | 117 | 16 |
|  | left paracingulate gyrus | 324 | 25 | - | - | - |
|  | left inferior temporal Gyrus posterior division | 302 | 30 | - | - | - |
|  | left temporal pole | 294 | 12 | - | - | - |
|  | left middle and inferior temporal gyrus posterior division | 289 | 21 | - | - | - |
|  | left middle and inferior temporal gyrus anterior division | 177 | 53 | - | - | - |
|  | left middle temporal gyrus anterior division | 147 | 33 | - | - | - |
|  | left inferior frontal gyrus (pars triangularis & opercularis) | 280 | 22/18 | - | - | - |

**Supplementary Table S3.**
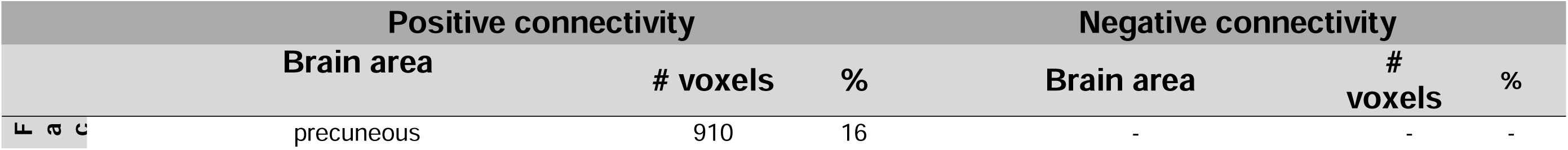
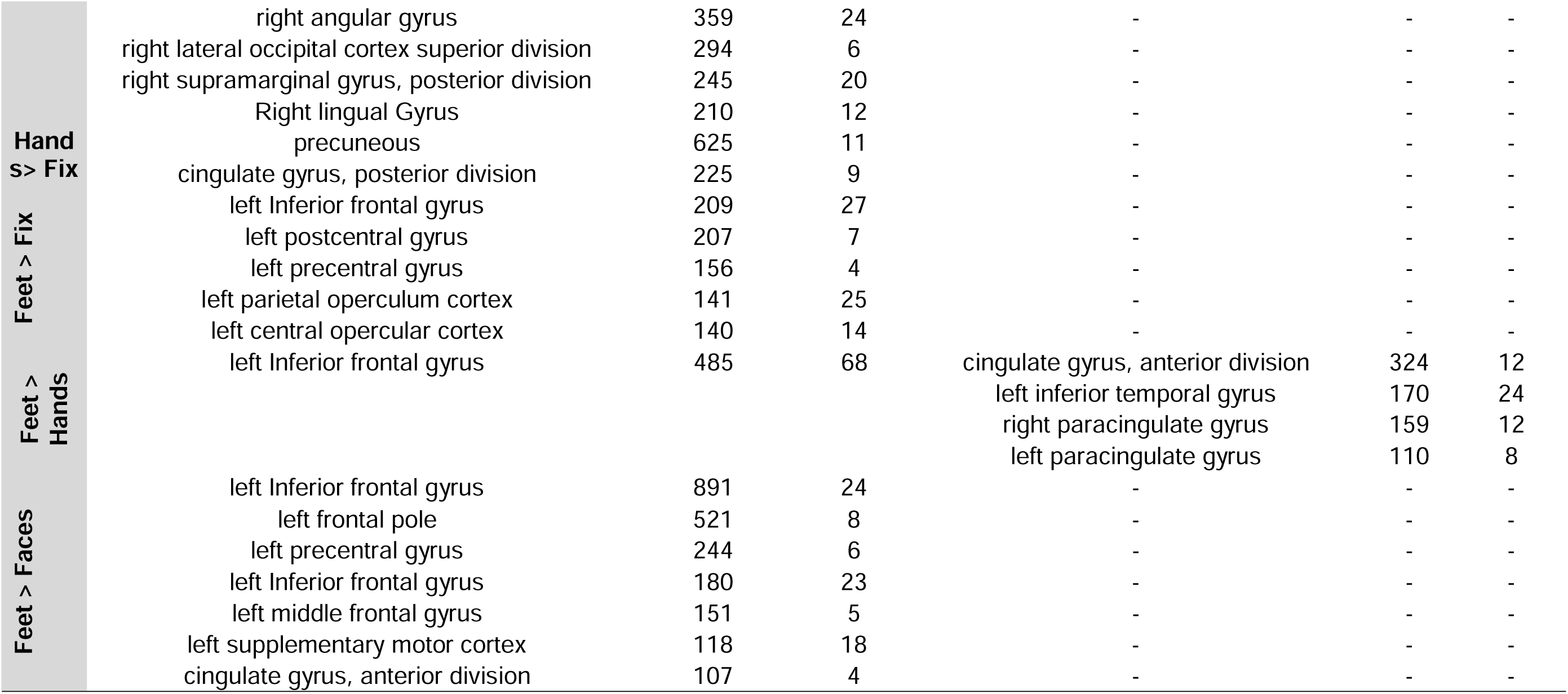
Connectivity results for Faces>Fixation, Hands>Fixation, Feet>Fixation, Feet>Hands and Feet>Faces for Tactile localizer contrast, left seed. Positive and negative brain connectivity for selected seed, number of voxels and % of areas activated.

